# Increased anxiety and decreased sociability in adulthood following paternal deprivation involve oxytocin in the mPFC

**DOI:** 10.1101/175380

**Authors:** Zhixiong He, Limin Wang, luo Luo, Rui Jia, Wei Yuan, Wenjuan Hou, Jinfeng Yang, Yang Yang, Fadao Tai

**Affiliations:** Institute of Brain and Behavioral Sciences, College of Life Sciences, Shaanxi Normal University, Xi’an, 710062, China; Cognition Neuroscience and Learning Division, Key Laboratory of Modern Teaching Technology, Ministry of Education, Shaanxi Normal University, Xi’an, 710062, China

**Author notes:** Correspondence to Fadao Tai, Institute of Brain and Behavioral Sciences, College of Life Sciences, Shaanxi Normal University, Xi’an, Shaanxi 710062, China.

## Abstract

Early adverse experiences often have devastating consequences on adult emotional and social behavior. However, whether paternal deprivation (PD) during the pre-weaning period affects brain and behavioral development remains unexplored in socially mandarin vole (*Microtus mandarinus*). We found that PD increased anxiety-like behavior and attenuated social preference in adult males and females; decreased prelimbic cortex OT-immunoreactive fibers and paraventricular nucleus OT positive neurons; reduced levels of medial prefrontal cortex (mPFC) OT receptor protein in females and OT receptor and V1a receptor protein in males. Intra-prelimbic cortical OT injections reversed anxiety-like behavior and social preferences affected by PD, whereas injections of OT and OT receptor antagonist blocked this reversal. These findings demonstrate that PD leads to increased anxiety-like behavior and attenuated social preferences with involvement of the mPFC OT system. The prelimbic cortex OT system may be an important target for the treatment of disorders related to early adverse experiences.

## Introduction

Social attachments are necessary in many species as they facilitate reproduction, increase survival, provide a sense of security and reduce feelings of stress and anxiety (Coria-Avila et al., 2014). In humans, attachment is especially important during early development because disruption of filial attachment in children (e.g., abuse, neglect, death of a parent) increases their vulnerability to mood and anxiety disorders at a later age (Brown et al., 1977; Agid et al., 1999; Bernet and Stein, 1999; Reinherz et al., 1999).

Neonatal social or paternal deprivation (PD) has been proven to exert a profound and persistent influence on the physiological and behavioral development of offspring. Neonatal rodents depend on their parents physiologically and emotionally. For example, PD impairs sociability (Bambico et al., 2013; Jia et al., 2009) and social recognition (Cao et al., 2014), reduces parental behaviors (Jia et al., 2009) and alloparental behavior (Ahern and Young, 2009), and inhibits the formation of pair bonding (Yu et al., 2013). A lack of paternal care can also affect emotional behavior (Ovtscharoff, 2006; Jia et al., 2009). In humans, PD impairs psychological and mental development, and increases the risk for substance abuse and personality disorders (Grossmann et al., 2002; Jablonska and Lindberg, 2007; Sobrinho et al., 2012). Despite emerging evidence of the impact of PD, the primary focus has been on father-offspring relationships from postnatal day (PND) 0–21 in rodents (Helmeke et al., 2009; Gos et al.,2014; Ahern and Young, 2009; Jia et al., 2011; Yu et al., 2015).

Mandarin voles (*Microtus mandarinus*) are furred by around PND 7, suggesting that their thermoregulatory ability is well developed. They open their eyes and begin to eat solid food around PND 13. Paternal care (e.g., licking, retrieving nest building) gradually declines from PND 14–20 (Wang, unpublished data). Pups from PND 1–13 and 14–21 should have different needs and the effects of disruption of father-offspring bonds during the latter period may not be the result of disrupted direct care, but the result of disrupted emotional attachment (He et al., 2017). This species is socially monogamous and exhibits extensive biparental investment and high offspring survival and growth (Tai et al., 2001; Tai and Wang, 2001). Mandarin vole pups have high levels of attachment to their fathers from PND 14–21 (He et al., 2017). Mandarin voles are an ideal model to investigate the effects of disruption of father-pup attachment on the brain and behaviors at adulthood because paternal care and biparental rearing patterns are found only in a minority of mammalian species (Kleiman and Malcolm, 1981). Whether PD from PND 14–21 affects emotional and social behavior remains unexplored.

The neuropeptide oxytocin (OT) is produced primarily in neurons of the hypothalamic paraventricular nucleus (PVN) and supraoptic nucleus (SON) (Onaka, 2004). OT is strongly implicated in prosocial behavior (Marlin et al., 2015; Young and Wang, 2004; Burkett et al., 2016; Oettl et al., 2016; Ma et al., 2016; Wircer et al., 2017) and decreased anxiety-related behavior (Windle et al., 1997; Blume, 2008; Sabihi, 2014b). Previous studies have found that PD alters levels of OT receptor (OTR) mRNA expression in the brain (Cao et al., 2014), and neonatal OT-treatments have long-term effects on behavior and physiology in mandarin voles (Jia et al., 2009). OT binds to OTRs (Burkett et al., 2016) or vasopressin 1a receptors (V1aR) (Song et al., 2014) to affect social behavior. If pre-weaning PD changes social preference and emotion, we predict that it should also affect levels of OT and OTR.

OTRs and V1aRs are found in the medial prefrontal cortex (mPFC) (Smeltzer et al., 2006; Lieberwirth and Wang, 2016). Several studies suggest that the mPFC is critical for the expression of anxiety-like behavior (Shah and Treit, 2003; Lisboa et al., 2010; Saitoh et al., 2014; Wang et al., 2015) and social behavior (Sabihi et al., 2014a; Niu et al., 2017; Lee et al., 2016). This brain region is a heterogeneous cortical structure composed of subregions, including the anterior cingulate (Cg), prelimbic cortex (PLC) and infralimbic cortex (Heidbreder and Groenewegen, 2003). Several studies have shown involvement of the PLC in the regulation of anxiety (Sabihi et al., 2014b; Wang et al., 2015) and social behavior (Young et al., 2001; Carrier and Kabbaj, 2012). A recent report found that a subset of mPFC neurons elevate discharge rates when approaching a strange mouse but not when approaching non-social objects (Lee et al., 2016) indicating involvement of the mPFC in social behavior. However, whether OT in the PLC is involved in the manifestations of pre-weaning deprivation on emotion and social preference remains unclear.

Using socially monogamous mandarin voles we investigated the effects of PD from PND 14–21 on emotion and social preference, and levels of OT and OTR in specific brain regions. We then tested whether microinjection of OT into the mPFC can recover the effects of pre-weaning PD. We hypothesized that disruption of early emotional attachment between pups and fathers affects anxiety-like behavior and social preference in mandarin voles at adulthood, and that the OT system is likely involved in this process.

## Results

### Experiment 1: Effect of pre-weaning PD on anxiety-like behavior and social preference

It has previously been shown that offspring who experience neonatal maternal separation or early deprivation display high levels of anxiety-like behavior (Lee et al., 2007; Sachs et al., 2013; Wei et al., 2010; Koe et al., 2016; Rees et al., 2005). Neonatal adversity has been shown to induce changes in social behavior, including avoidance, fear and decreased social interaction (Giachino et al., 2007; Jia et al., 2009; Toth et al., 2013). Experiment 1 was designed to test the hypothesis that pre-weaning PD (PND 14–21) increases anxiety-like behavior and reduces social preference. At 70 days of age, subjects were randomly divided into the control group (PC group: 7 females and 7 males) and PD group (7 females and 7 males).

An open field test (OFT) showed that the percentage of time spent in the central area was greater in PC group compared with the PD group (male: t(12) = 4.158, p < 0.01; female: t(12) = 3.226, p < 0.05). However, the total distance covered in the PC was not different from the PD group (Fig. 1).

**Fig 1.**
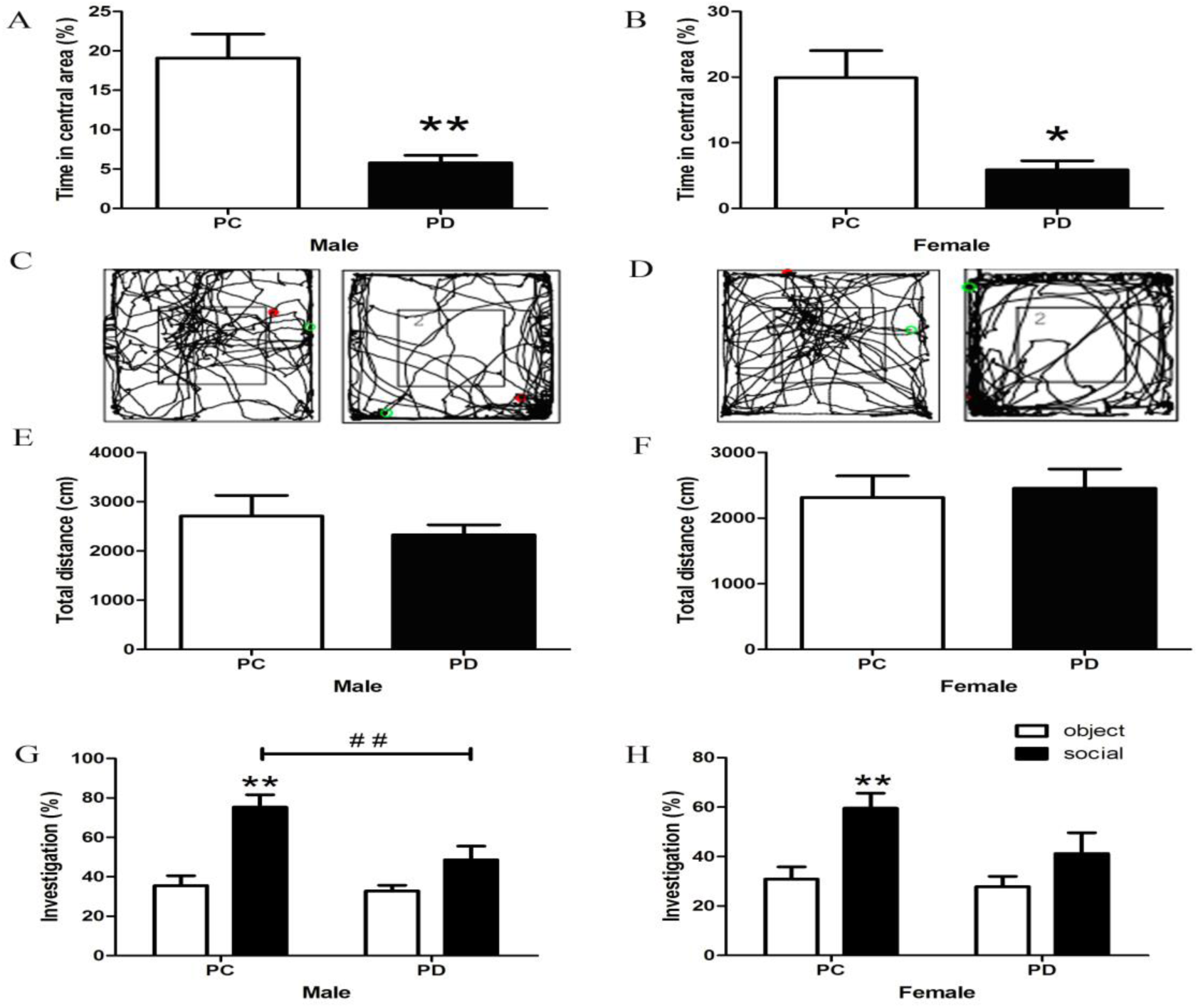
Effect of PD on anxiety-like behavior and social preference in adult mandarin voles. (**A, B**) Percentage of time in the central area, (**C, D**) representative path and (**E, F**) total distance of mandarin voles in the open field test (Male: PC: n = 7, PD: n = 7; Female: PC: n = 7, PD: n = 7). *p < 0.05; **p < 0.01. Independent sample t-tests. Effect of PD on social preference in (**G**) male and (**H**) female mandarin voles (Male: PC: n = 7, PD: n = 7; Female: PC: n = 7, PD: n = 7). Error bars indicate SEM. **p < 0.025 vs. object stimulus. ## p < 0.025 vs. PC. Two-way ANOVA (factors: treatment × stimulus). PC: biparental care; PD: paternal deprivation.

Two-way ANOVA indicated a significant interaction between the percentage of investigation time in males for treatment x absence/presence of a stimulus mandarin vole interaction (F(1,24) = 4.775, p < 0.05). The PC group engaged in more investigation of the social stimulus than object stimulus in males (p < 0.01). Pre-weaning PD caused decreased social investigation in males during the social preference test (p < 0.01). A main effect of absence/presence of a stimulus mandarin vole was found (F(1,24) = 11.785, p < 0.01), such that PC group females engaged in more investigation of the social stimulus than object stimulus (p < 0.01). PD treatment did not produce a main effect (F(1,24) = 3.064, P = 0.093) in females. There was also no significant interaction in females (F(1,24) = 1.553, P = 0.225). PD did not affect investigation of the social stimulus and object stimulus in males or females (Fig. 1).

### Experiment 2: Effect of pre-weaning PD on mPFC and PVN OT-IR

Previous studies have shown that neonatal isolation or maternal separation results in a decrease in OT-IR neurons in the PVN in male adult mandarin voles and female mice (Wei et al., 2013; Veenema et al., 2007). It is apparent that OT production in the hypothalamus is altered in response to social interactions in many species (Leng et al., 2008). Tactile stimulation at an early developmental stage induces immediate-early gene activity in OT neurons in prairie voles (Barrett et al., 2015) and rabbit pups (Caba et al., 2003), lower stress responses during their adulthood (Liu et al., 1997; Meaney et al., 2001), and alleviates the negative effects of neonatal isolation on novel object recognition and sociability (Wei et al., 2013). Oxytocinergic neurons within the PVN project to the mPFC (Uvnäs-Moberg et al., 2015) and the NAcc (Ross et al., 2009). We therefore tested levels of OT-IR in the NAcc, mPFC and PVN. Three hours after the social preference test, experimental mandarin voles from the PC (n = 4 males, 4 females) and PD (n = 4 males, 4 females) groups were anesthetized with pentobarbital sodium. Brains were removed from the skull and placed in 4% paraformaldehyde three days. Brains were then dehydrated in 30% sucrose solution until saturated at 4°C for immunohistochemistry.

Two-way ANOVA revealed interactions between treatment x sex for PVN OT-IR neurons (F(1,12) = 8.265, p < 0.05) and mPFC OT-IR fibers (F(1,12) = 17.621, p < 0.01). Post-hoc tests indicated that PD group males and females had fewer PVN OT-IR neurons than the PC group (both p < 0.01) (Fig. 2), and reduced mPFC OT-IR fibers than PC group males and females (both p < 0.01) (Fig. 3). Females possessed more PVN OT-IR neurons (p < 0.01) and mPFC OT-IR fibers (p < 0.01) than males (Fig. 2, 3). NAcc OT-IR fibers did not differ between groups (Fig. 4).

**Fig 2.**
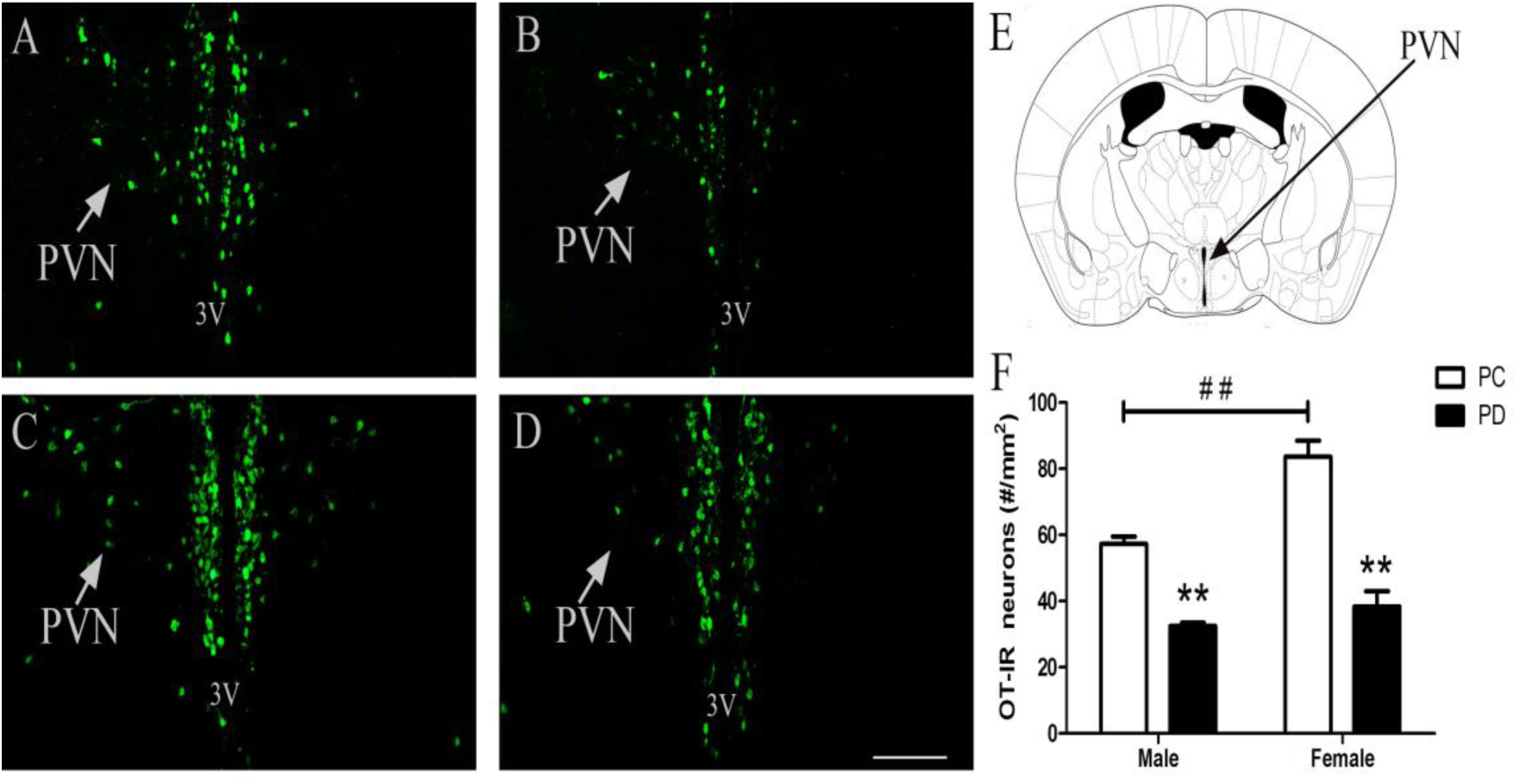
Effect of PD on PVN OT-ir neurons. (**A)** PC males; (**B**) PD males; (**C**) PC females; (**D**) PD females; magnification 10 x, scale bar 200 *μ*m, 3V–3rd ventricle. (**E**) Schematic drawing illustrating tissue in the PVN. (**F**) Quantification of OT-IR neurons in the PVN (Male: PC: n = 4, PD: n = 4; Female: PC: n = 4, PD: n = 4). Error bars indicate SEM. **p < 0.01 vs. PC. ## p < 0.01 vs. male. Two-way ANOVA (factors: treatment × sex). PVN: paraventricular nucleus.

**Fig 3.**
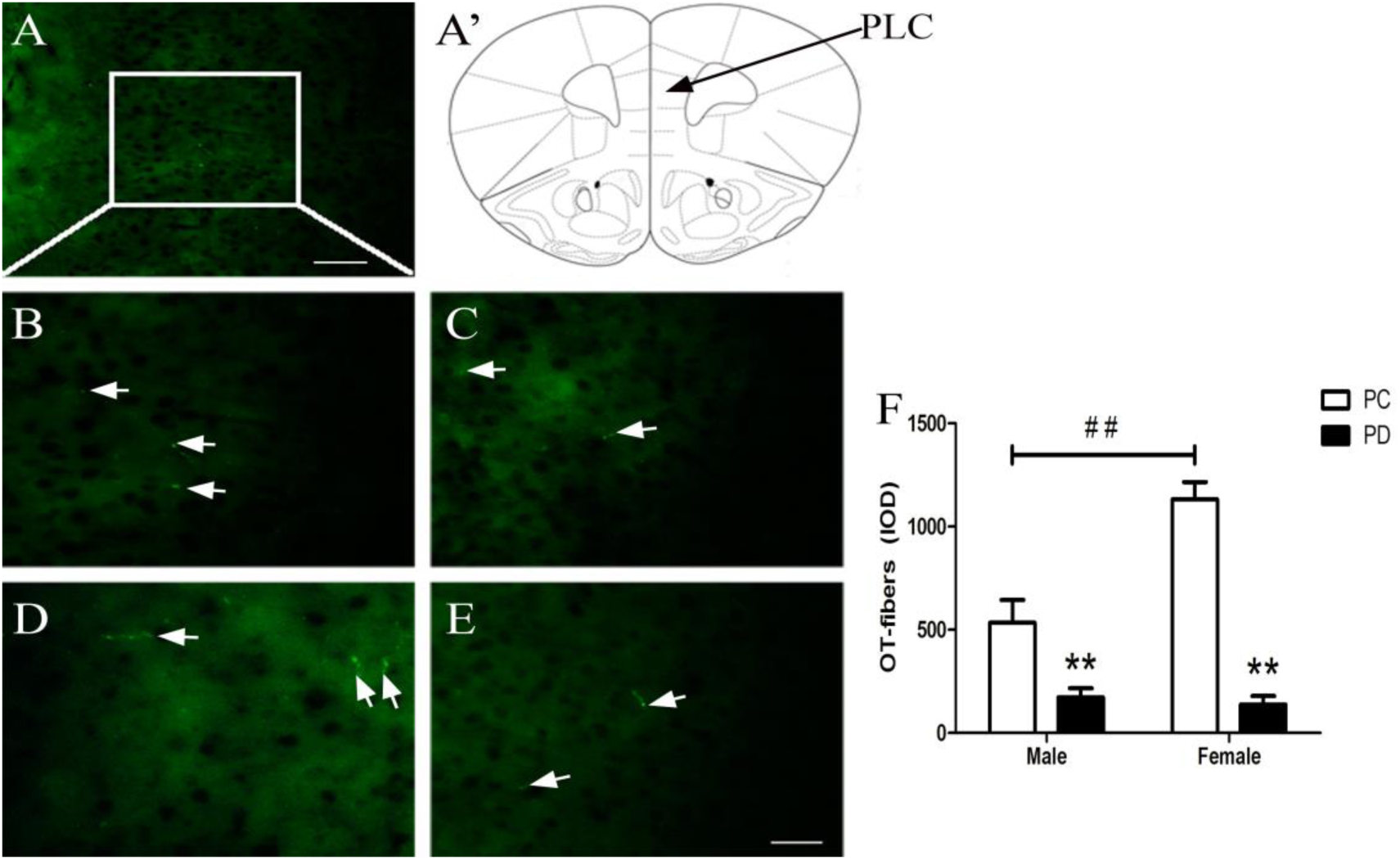
Effect of PD on PLC OT-ir fibers. (**A-A’**) Photomicrograph showing high somatodendritic immunoreactivity of OT in the PLC (magnification 20 x, scale bar 100 *μ*m). (**B)** PC males; (**C**) PD males; (**D**) PC females; (**E**) PD females; magnification 40 x, scale bar 50 *μ*m. (**F**) Quantification of OT-IR fibers in the PLC (Male: PC: n = 4, PD: n = 4; Female: PC: n = 4, PD: n = 4). Error bars indicate the SEM. ## p < 0.01 vs. male. Two-way ANOVA (factors: treatment × sex). PLC: prelimbic cortex.

**Fig 4.**
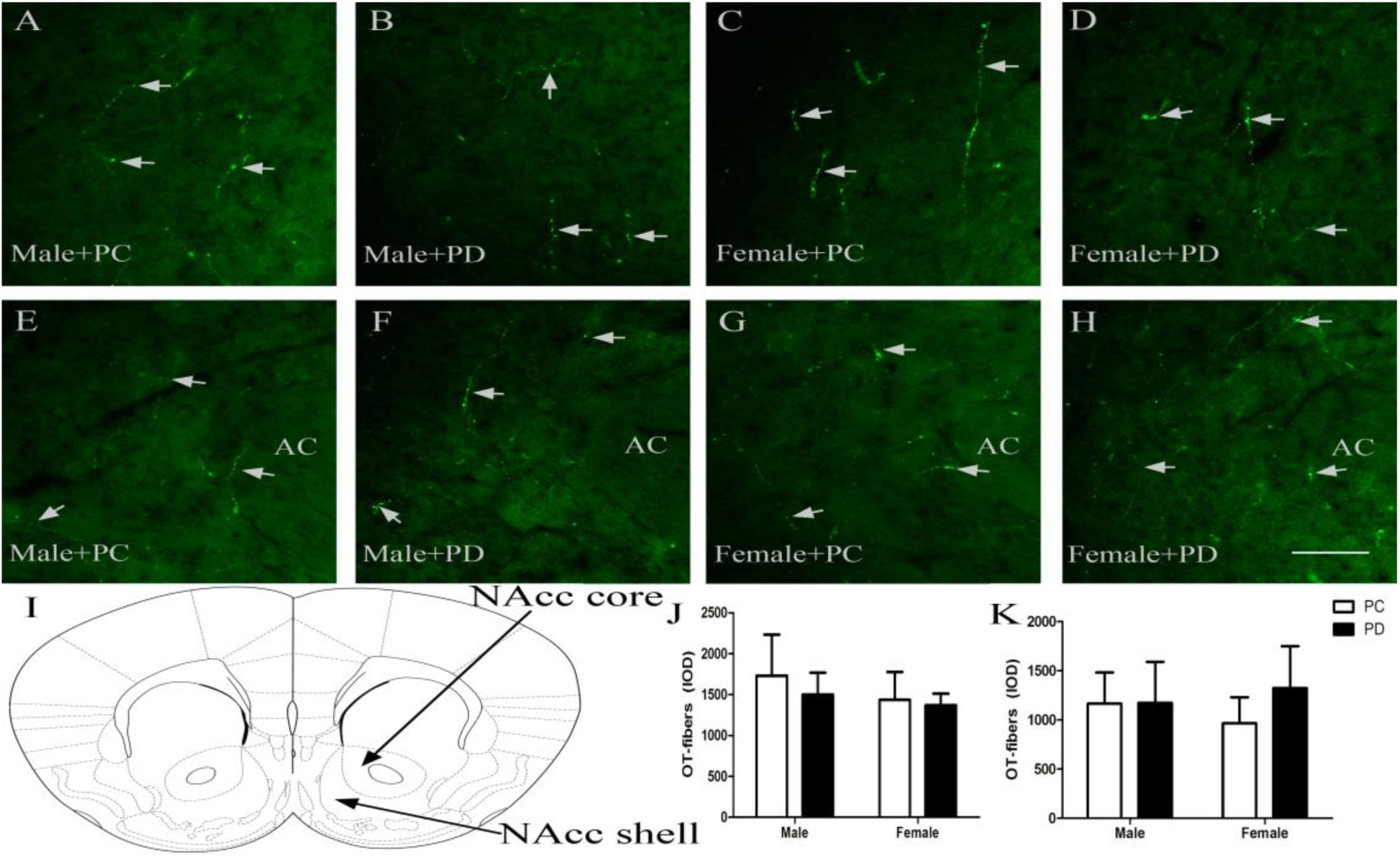
Effect of PD on NAcc OT-IR fibers in males and females. (**A-D**) NAcc shell; (**E-H**) NAcc core, AC = anterior commisure; magnification 20 x, scale bar 100 *μ*m. (**I**) Schematic drawing illustrating tissue in the NAcc. (**J-K**) Quantification of OT-IR fibers in the NAcc (Male: PC: n = 4, PD: n = 4; Female: PC: n = 4, PD: n = 4). Error bars indicate SEM. Two-way ANOVA (factors: treatment × sex). NAcc: nucleus accumbens.

### Experiment 3: Effect of pre-weaning PD on mPFC OTR-IR and V1aR-IR, serum OT and corticosterone (CORT) concentration

PD reduced OT-IR fibers in the PLC (Experiment 2). OT and arginine vasopressin (AVP), and OTR and V1aR, display a high degree of sequence homology and both peptides can activate both receptors (Chini and Manning, 2007). OTR (Li et al, 2016), V1aR and AVP (Dumais and Veenema, 2016) are involved in regulation of mood and social behavior. Experiment 3 was designed to test the hypothesis that pre-weaning PD alters OTR, V1aR and/or AVP densities in the mPFC, NAcc and/or PVN. Voles from the PC (n = 4 males, 4 females) and PD (n = 4 males, 4 females) groups were sacrificed by rapid decapitation at 70 days of age. All brains were harvested, frozen on dry ice, and stored at −80°C until Western blotting.

PD group males had lower levels of OTR (t(6) = 2.799, p < 0.05) and V1aR (t(6) = 2.816, p < 0.05) proteins in the mPFC than PC group males. No group differences were noted for levels of male OTR and V1aR protein in the NAcc or PVN. PD group females had lower levels of OTR protein in the mPFC than PC group females (t(6) = 2.648, p < 0.05), but not in the NAcc or PVN. No group differences were noted for levels of female V1aR protein in the mPFC, NAcc or PVN. PD had no effect on AVP neuropeptide in the mPFC, NAcc or PVN regardless of sex (Fig. 5).

**Fig 5.**
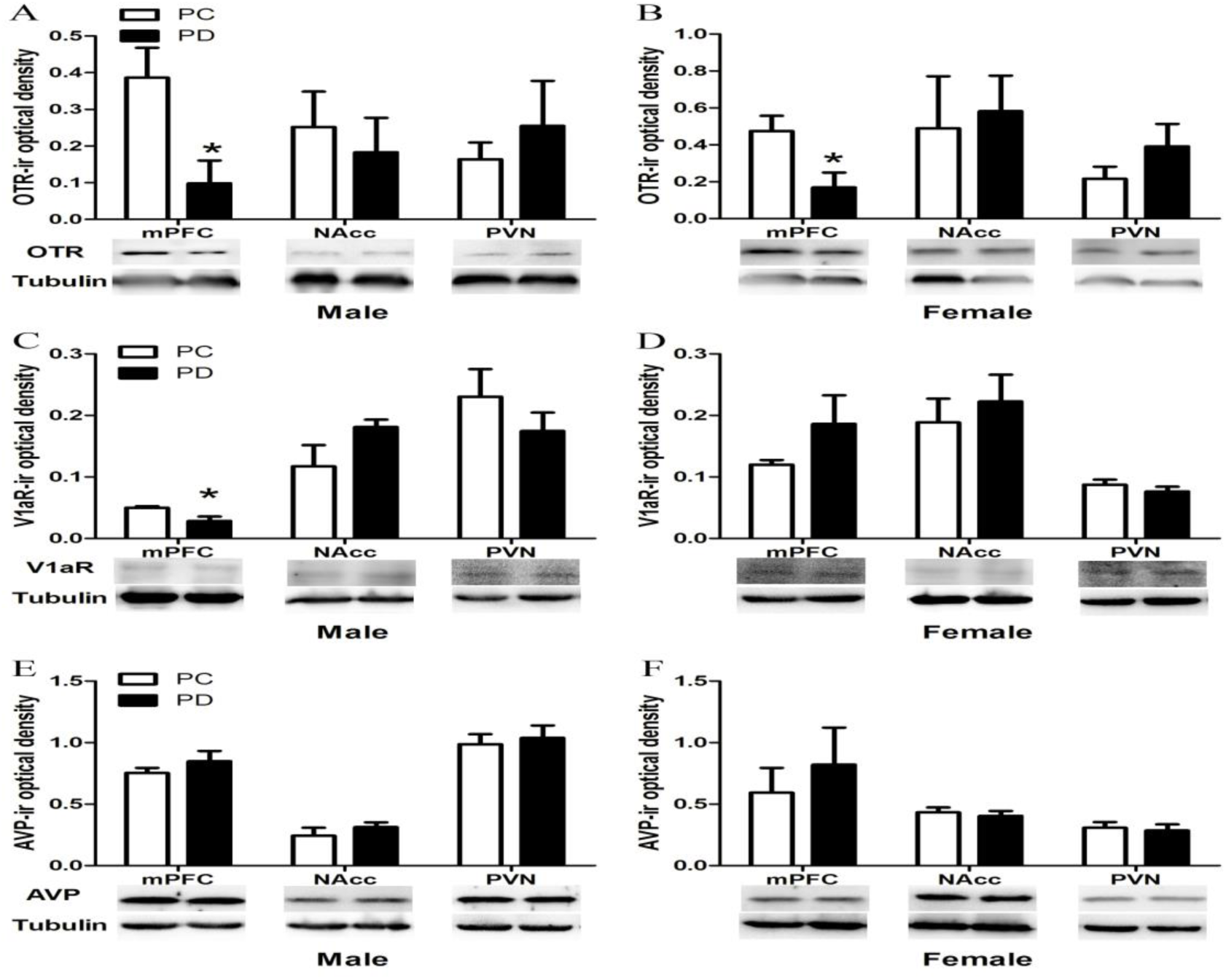
Effects of PD on mesocorticolimbic (**A-B**) OTR, (**C-D**) V1aR and (**E-F**) AVP immunoreactivity in male and female mandarin voles (Male: PC: n = 4, PD: n = 4; Female: PC: n = 4, PD: n = 4). Error bars indicate SEM. * p < 0.05. Independent sample t-tests. PC: biparental care; PD: paternal deprivation; OTR: oxytocin receptor; V1aR: vasopressin 1a receptor; AVP: arginine vasopressin; mPFC: medial prefrontal cortex; NAcc: nucleus accumbens; PVN: paraventricular nucleus.

PVN OT signaling is necessary and sufficient for social buffering effects (Smith and Wang, 2014) and is anxiolytic (Neumann et al., 2000) by suppressing hypothalamic-pituitary-adrenal (HPA) axis function. To assess whether PD reduces OT and increases CORT in serum, three hours following the social preference test, experimental mandarin voles from the PC (n = 6 males, 6 females) and PD (n = 6 males, 6 females) groups were anesthetized with pentobarbital sodium (30 mg/kg i.p.). Serum OT and CORT concentrations were monitored.

Two-way ANOVA revealed an interaction between treatment x sex for serum OT (F(1,20) = 5.055 p < 0.05) and CORT (F(1,20) = 11.755, p < 0.01). The post-hoc test indicated that pre-weaning PD significantly reduced serum OT (p < 0.01) and increased serum CORT (p < 0.01) concentrations only in females. Female serum OT levels were much higher than males in the PC group (p < 0.05). Female serum CORT concentrations were also much higher than males in the PD group (p < 0.01) (Fig. 6).

**Fig 6.**
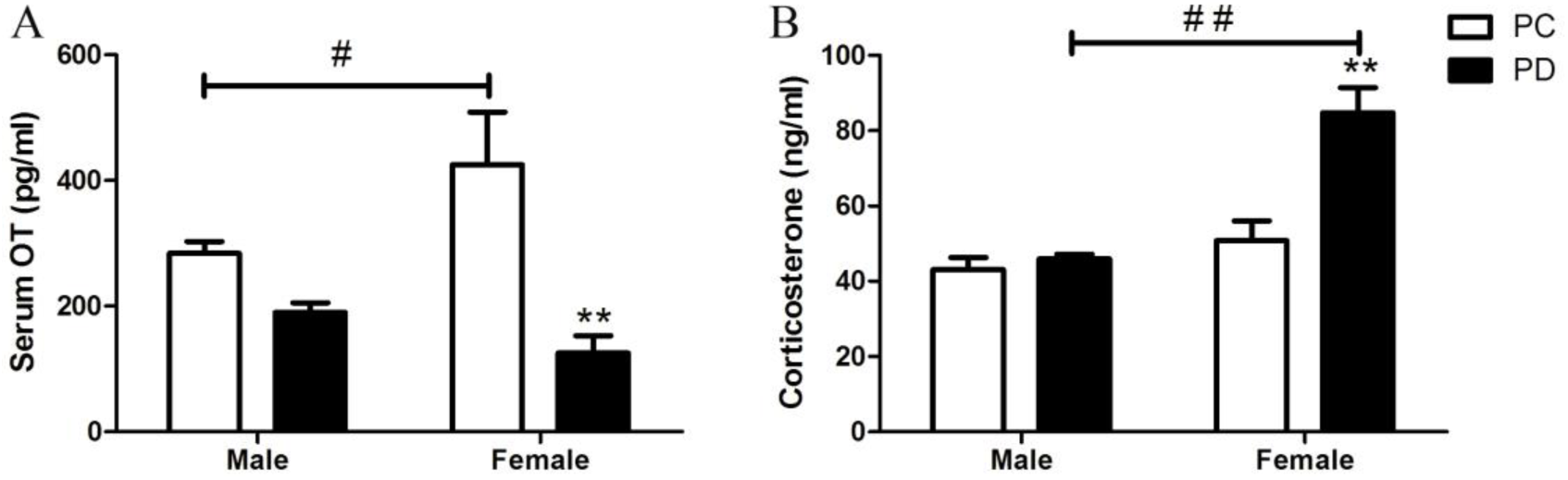
Levels of serum (**A**) OT and (**B**) CORT concentrations in adult mandarin voles (Male: PC: n = 6, PD: n = 6; Female: PC: n = 6, PD: n = 6). Error bars indicate the SEM. **p < 0.01, PC vs. PD. # p < 0.05, ## p < 0.01, vs. male. Two-way ANOVA (factors: treatment × sex). PC: biparental care; PD: paternal deprivation.

### Experiment 4: Effect of microinjection of OT into the PLC on anxiety-like behavior and social preference altered by pre-weaning PD

PD reduced OT-IR fibers and OTR protein density in the mPFC (Experiment 2, 3) and OT infused into the PLC region of the mPFC reduces anxiety-like behavior (Sabihi et al., 2014b). OTRs are important for modulation of social and emotional behavior in the mPFC (Li et al, 2016). Therefore, we tested the hypothesis that microinjection of OT in the PLC restores anxiety-like behavior and social preference altered by pre-weaning PD. To this end, we implanted bilateral injection cannulas into the PLC. After three days of recovery, subjects received an intra-PLC injection (Fig. 7A–C) of CSF (male = 6, female = 6), OT (male: 1 ng OT = 6, 10 ng OT = 6; female: 1 ng OT = 6, 10 ng OT = 6) or OT plus OTA (male: 10 ng OT/10 ng OTA = 6, 10 ng OT/100 ng OTA = 5; female: 1 ng OT/10 ng OTA = 5, 1 ng OT/100 ng OTA = 5). Levels of anxiety-like behavior and social preference were then measured.

**Fig 7.**
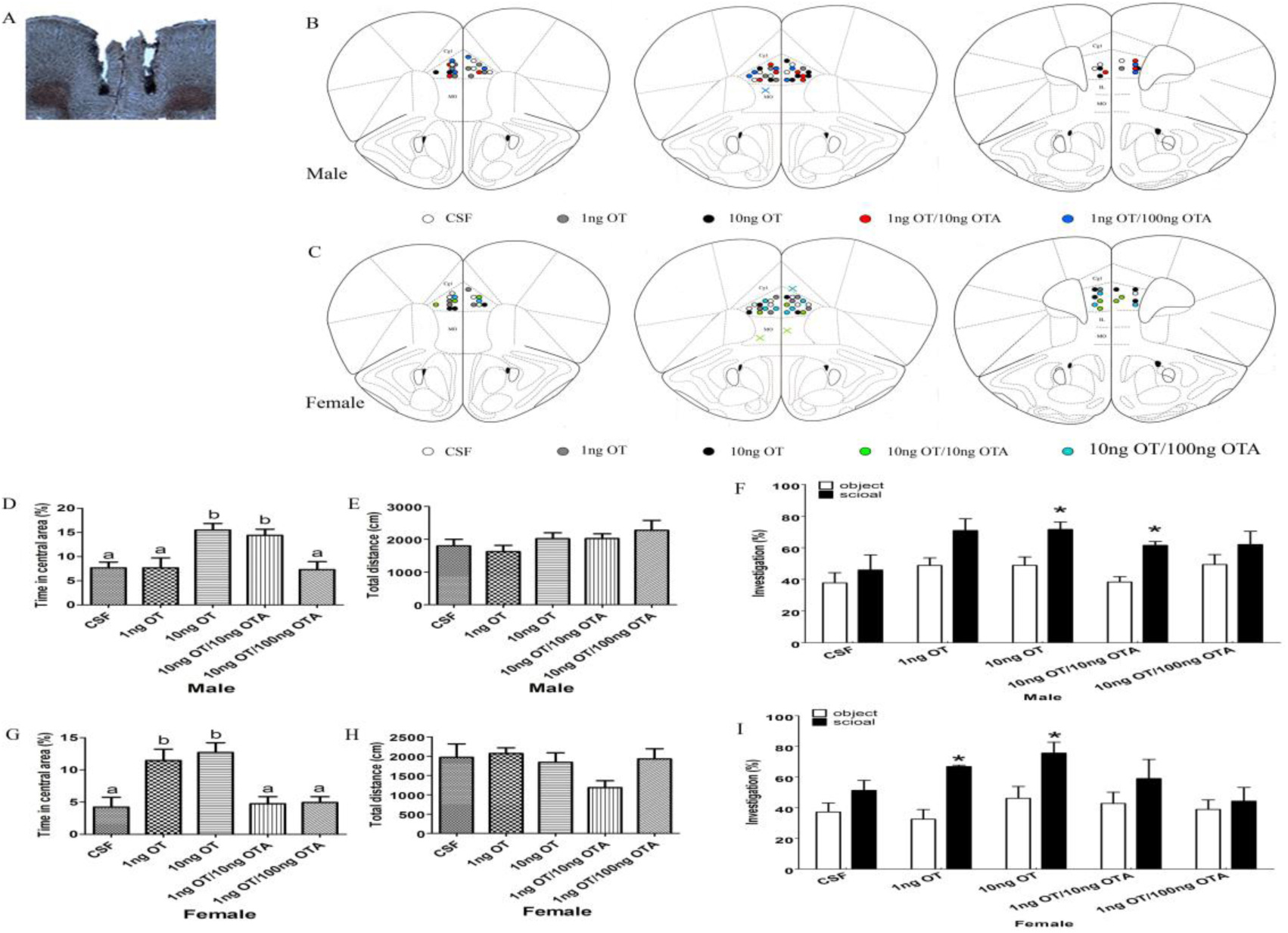
Effects of PLC OT administration on anxiety-like behavior and social preference following the disruption of early emotional attachment in adult mandarin voles. (**A**) Histological representations of microinjection site and (**B, C**) schematic diagrams showing the location of injector tips in the PLC. **×**: missed. OT in the PLC is anxiolytic in both of sexes. (**D, G**) Percentage of time in the central area and (**E, H**) total distance in the open field test (Male: CSF: n = 6, 1ng OT: n = 6, 10ng OT: n = 6, 10ng OT /10 ng OTA: n = 6, 10ng OT /100 ng OTA: n = 5; Female: CSF: n = 6, 1ng OT: n = 6, 10ng OT: n = 6, 1ng OT /10 ng OTA: n = 5, 1ng OT /100 ng OTA: n = 5). Bars without the same letters are significantly different. One-way ANOVA. OT in the PLC promotes a social preference in (**F**) males (CSF: n = 6, 1ng OT: n = 6, 10ng OT: n = 6, 10ng OT /10 ng OTA: n = 6, 10ng OT /100 ng OTA: n = 5) and (**I**) females (CSF: n = 6, 1ng OT: n = 6, 10ng OT: n = 6, 1ng OT /10 ng OTA: n = 5, 1ng OT /100 ng OTA: n = 5). * p < 0.01 vs object stimulus. Two-way ANOVA (factors: treatment × sex).

One-way ANOVA showed that microinjection of OT in the PLC increased the percentage of time spent in the central area for males (10 ng: p < 0.01) and females (1 ng: p < 0.05 and 10 ng: p < 0.01) exposed to pre-weaning PD, while OT plus either dose of OTA had no effect (except male 10 ng OT/10 ng OTA, p < 0.05). No differences were found between treatment groups for total distance travelled (Fig. 7).

Two-way ANOVA found a main effect of the absence/presence of a stimulus mandarin vole in both sexes (male: F (1,48) = 23.254, p < 0.01; female: F(1,46) = 18.952, p < 0.01). However, there were no interactions between treatment and the absence/presence of the stimulus vole in both sexes. OT-treated males (10 ng) and females (1 ng and 10 ng) demonstrated a preference for the social stimulus over the object stimulus (p < 0.01), while subjects treated with OT plus either dose of OTA showed no preference (except for males receiving 10 ng OT/10 ng OTA, p < 0.01) (Fig. 7).

## Discussion

The present study found that pre-weaning PD possibly disrupts emotional attachment in mandarin vole pups, as evidenced by increased levels of anxiety-like behavior and attenuated social preference in male and female adults (Experiment 1). PD reduced mPFC OT-IR fibers and PVN OT-IR neurons (Experiment 2), and decreased mPFC OTR protein in females, OTR and V1aR protein in males, and reduced serum OT and increased CORT in females (Experiment 3). We then demonstrated that intra-PLC OT injection restored anxiety-like behavior and social preference altered by pre-weaning PD (Experiment 4).

In this study, PND 14–21 PD increased anxiety-like behavior and reduced social preference in both sexes. This is consistent with studies in animals and other humans that early severe deprivation is associated with behavioral abnormalities (Rosenblum and Harlow, 1963; Rutter et al., 2001). The current result is also supported by one of our previous studies that PND 14–21 offspring show emotional attachment to fathers (He et al., 2017). Thus, PND 14–21 PD should disrupt emotional attachment to fathers and adversely affect emotion and sociability. These effects are similar to previous findings that neonatal PD during PND 1–21 increases anxiety (Yu et al., 2011) and reduces sociability (Jia et al., 2009; Bambico et al., 2013; Farrell et al., 2016). We infer that the disruption of attachment between pups and fathers during PND 14–21 increases levels of anxiety and reduce levels of sociability.

The present study found that PND 14–21 PD reduced mPFC OT-IR fibers and OTR protein in both sexes, but did not affect levels of AVP. The PLC may receive long range axonal projections from OT-producing neurons. PD also decreased OT-IR neurons in the PVN. Because oxytocinergic neurons within the PVN project to the mPFC (Lieberwirth and Wang, 2016), a decrease in OT-IR neurons in the PVN possibly led to a reduction in OT-IR fibers in the mPFC. A previous study found that OT in the PL region of the mPFC decreased anxiety regardless of sex, and neither AVP nor OTR-A affected anxiety-like behavior (Sabihi et al., 2014b). Blocking OTR in the mPFC enhances postpartum anxiety, but has no effect on anxiety in virgin females (Sabihi et al., 2014a). OTR knockout mice display deficits in social approach behavior (Nishimori et al., 2008) and social memory (Lee et al., 2008). Thus, OTR in the mPFC may play different roles under different physiological and pathological conditions. PVN OT neurons predominantly express vesicular glutamate transporter 2, suggesting that depolarization of these neurons is coupled with synaptic glutamate release in their projections such as those to the mPFC (Kawasaki et al., 2005; Johnson and Young, 2017). In addition, a specific class of interneurons (OxtrINs) in the mPFC is critical to the modulation of social and emotional behavior in both sexes (Li et al, 2016). It is probable that OT neurons releasing Glutamate activate local GABA-interneurons in the mPFC, leading to reduced anxiety and regulated social behavior. Thus, we conclude that disruption to attachment between pups and fathers increases levels of anxiety and impairs social preference via a decrease in OT-IR fibers and OTR protein in the mPFC.

PND 14–21 PD decreased V1aR protein in males but had no effect on levels of V1aR protein in females. Sex differences in the OT system may therefore be implicated in sex-specific regulation of impaired social behavior (Smeltzer, 2006; Dumais and Veenema, 2016). V1aR and OTR show distinct and largely nonoverlapping expression in the rodent brain (Dumais and Veenema, 2016). Female prairie voles had higher densities of OTR binding but lower densities of V1aR binding than males in the mPFC (Smeltzer, 2006). Intracerebroventricular (ICV) injections of OT induce social communication by activating V1aR in male Syrian hamsters (Song et al., 2014). Studies have shown that some of the prosocial effects of OT may be mediated by the V1aR in males (Sala et al., 2011; Ramos et al., 2013) and that V1aR knockout mice have impaired social interaction (Egashira et al, 2007). V1aR is G protein-coupled receptor, similar to the OTR. We speculate that OT possibly regulates anxiety-like behavior and social preference in males via OTR and V1aR in the PLC.

PND 14–21 PD did not affect NAcc OT-IR fibers and OTR-IR densities in either sex. This is inconsistent with the previous finding that PD (PND 1–21) alters levels of OTR mRNA expression in the NAcc (Cao et al., 2014). These discrepancies may result from different deprivation periods, however, this requires further experimentation.

PND 14–21 PD reduced serum OT levels and increased serum CORT levels in females, but not males. This result may be consistent with our previous work (Cao et al., 2014; Wu et al., 2014). We also found that disruption of early emotional attachment reduced OT-IR neurons in the PVN. These results suggest that OT magnocellular neurons in the hypothalamic nuclei may have reduced OT into the peripheral circulation via the posterior pituitary (Ludwig and Leng, 2006). During some prosocial behavior, OT is released into plasma and centrally in females (Ross et al., 2009; Churchland and Winkielman, 2012). Maternal separation is known to decrease PVN OT-IR neurons in females, but not males (Veenema et al., 2007). One possibility is that females have higher ratings of fear, irritability and unhappiness under psychosocial stress compared to males (Lee et al., 2013). Another reason for these discrepancies may be different treatments (maternal separation and PD) and different sensitivities to neonatal stress in different species. OT can directly modulate the stress reactive HPA axis (Acevedo-Rodriguez, et al., 2015). Females exhibit not only higher HPA reactivity under basal conditions than males, but also after chronic stress (Hillerer et al., 2013). ICV (Windle et al., 1997) and PVN (Smith and Wang, 2014) administration of OT decreases circulating CORT and anxiety-like behavior. Thus, higher serum CORT induced by pre-weaning PD may be explained in part by the decrease in PVN OT.

A major finding in the current study is that OT microinjection directly into the PLC of males (10 ng) and females (1 ng and 10 ng) restored changes to anxiety-like behavior and social preference resulting from PD, while voles treated with OT plus either dose of OTA did not exhibit reversal of any kind (except for the male 10 ng OT/10 ng OTA group). This is consistent with previous findings that OT in the PLC reduces anxiety-like behavior in both sexes (Sabihi et al., 2014b). Injection of highly specific OTA into the PLC region of the mPFC increases anxiety-like behavior in postpartum females (Sabihi et al., 2014a) and OTR blockade in the postpartum PLC impairs maternal care behavior and enhanced maternal aggression (Sabihi et al., 2014a). OxtrINs were identified in the mouse mPFC that express OTR and are activated in response to OT (Nakajima et al, 2014). OxtrINs are important for the modulation of social and emotional behavior in males and females, and are a molecular mechanism that acts on local mPFC circuits to coordinate responses to OT and corticotropin-releasing hormone (Li et al, 2016). Similarly, OT infusion into the PLC reverses amphetamine-induced deficits in social bonding (Young et al., 2014). OT and dopamine (DA) interaction regulate affiliative social behaviors (Liu and Wang, 2003; Shahrokh et al., 2010). Father absence in the monogamous California mouse impairs social behavior and decreases pyramidal neuronal responses to DA in the mPFC (Bambico et al., 2013). Oxytocin injected into the ventral tegmental area increased extracellular DA concentration in the dialysate from the mPFC (Sanna et al., 2012). Therefore, there is likely to be a reciprocal interaction between OT and DA to regulate social behavior in the mPFC, and mPFC OT may be a promising target for treating emotional and social disorders induced by adverse early experiences.

Mandarin voles can be used as an animal model to investigate the effects of early emotional attachment disruption on the adult brain and behavior and underlying mechanism (He et al., 2017). Disruption to early emotional attachment between pups and fathers impairs emotional and social behavior and leads to OT system dysfunction in the brain. We provide intriguing evidence that site-specific OT action in the PLC has potential beneficial effects on the recovery of emotional and social dysfunction induced by the disruption to early emotional attachment. The modulation of OT on emotion and social behavior was sex-specific. Therefore, OT may be potentially targeted to ameliorate social and emotional deficits resulting from early adverse experiences.

## Methods and Materials

### Subjects

Animals were a laboratory-reared offspring originating from a wild population of mandarin voles in Henan, China. Animals were maintained on a 12:12 light: dark cycle with unlimited access to food (carrot and rabbit chow) and were provided with water and cotton nesting material in polycarbonate cages (44 cm x 22 cm x 16 cm).

For the paternal deprivation (PD) treatment, fathers (F1 generation) were removed permanently from the home cage after pups (F2 generation) were 14 days of age until weaning at PND 21. For the biparental care control group (PC), all family members were housed in their home cage and left undisturbed until pups were weaned at 21 days of age. Offspring at 70 days of age were tested using the behavioral paradigms below. In the female, only experimental data from diestrous individuals were included to avoid effects from the estrous cycle.

### Open field test (OFT)

To assess the impact of pre-weaning PD on adult anxiety-like behavior, F2 mandarin voles were observed in the OFT on PND 70. Subjects were placed in a center of an open-field arena (50 cm x 50 cm x 25 cm), and the duration and distance moved within the center or periphery was recorded using an automated system (SocialScan 2.0, Clever Sys, Reston, VA, USA). Measures include the proportion of time spent in the central area and total distance moved in the OFT.

### Social preference test (SPT)

Immediately after the OFT, the SPT was carried out. The SPT was based on the social approach-avoidance paradigm previously described (Qiao et al., 2014). Briefly, prior to testing voles were placed in a box. The test box (50 cm length x 50 cm width x 24 cm height) was constructed of white glacial polyvinylchloride. After 5 min of habituation, an empty wire-mesh cage (object stimulus; 10 cm length x 10 cm width) was placed near one side wall of the arena for 10 min, which was then exchanged for a cage containing an stranger same-sex con-specific (social stimulus) for an additional 10 min. Behavioral responses to an empty cage or to a cage with a stimulus individual were videotaped, and scored and quantified afterwards using OBSERVER v5.0 (vNoldus, NL). Measures included the time spent investigating the object and social stimulus. Data are presented as investigation time/total time (10 min) x 100%.

### Elisa

Blood was collected directly in microcentrifuge vials from the heart (Cao et al., 2014). After clotting, blood was centrifuged at 6000 rpm for 30 min at 4°C. Supernatant was collected. Serum OT and CORT levels were monitored by a vole-specific enzyme-linked immunosorbent assay (Shanghai Xitang Biotechnology, Shanghai, China), according to the manufacturers’ instructions. The resultant absorbance was measured at 450 nm using a Metertech microplate reader (BioTek Instruments, Winooski, USA) after the reader was zeroed using the blank well. Variation between duplicate measurements was less than 5%.

### Immunohistochemistry

20*μ*m (mPFC) and 40 *μ*m (NAcc and PVN) thick coronal slices were prepared on a cryostat (CM1950, Leica, Germany). Sections were rinsed with 0.01 M phosphate buffer solution (PBS) for 10 minutes following incubation with 0.6% H_2_O_2_ for 20 min, and rinsed for 3 x 5 min with 0.01 M PBS. Sections were then preincubated for 60 min with in blocking solution (normal goat serum, AR0009, Boster Company). Sections were incubated for 48 h at 4 ^o^C in mouse monoclonal antibody OT (MAB5296, 1:7500, Chemicon-Millipore) diluted in antibody diluent (0.01 M PBS containing 20% bovine serum albumin and 1.7% Triton X-100). The following day, sections were rinsed 3 x 5 min with 0.01 M PBS and incubated in the secondary antibody DyLight 488-conjugated goat anti-mouse (BA1126, 1:200, Boster Company) for 60 min in a 37°C water bath. Afterwards, sections were rinsed for 3 x 5 min with 0.01 M PBS and fixed with antifade solution (AR1109, Boster Company). Slices were photographed with a microscope and Nikon camera (Tokyo, Japan). For each vole, OT-immunoreactive fibers (mPFC and NAcc) and the number of OT-IR neurons (PVN) were counted, three representative sections from anterior to posterior anatomically matched between subjects were chosen to minimize variability. The count of OT-IR cells (Song et al., 2010) and OT-immunoreactive fibers (DiBenedictis et al., 2017) followed the method of the previous report. All immunohistochemistry procedures included sections of negative controls (the primary antibody was not added). An observer blind to experimental conditions performed the entire analysis. The number of positive neurons (PVN) and positive fibers (NAcc and mPFC) was quantified from images acquired under a 10X (PVN), 20X (NAcc) and 40X (mPFC) objective using the cell counter (PVN) and the value of integrated optical density (IOD) plugin in Image-pro-plus; these were averaged across three nonoverlapping sections in an evenly spaced series per animal.

### Western blotting

Brains were immediately extracted and frozen in dry ice. Coronal sections (200 μm) w ere cut on a cryostat and frost mounted onto microscope slides. Bilateral tissue punches with a 1 mm diameter were taken from the entire mPFC (Cg and PLC), NAcc and PVN and stored at −80°C until processing. Total proteins were extracted with RIPA lysis buffer containing protease inhibitors (R0010, Solarbio Biotechnology). Protein samples were separated by sodium dodecyl sulphate-polyacrylamide gel electrophoresis and transferred to PVDF membranes (Millipore, Billerica, MA, USA). The membrane was incubated with the following diluted primary antibodies: OTR (ab181077, 1:2000, Abcam), V1aR (GTX89114, 1:7000, GeneTex), AVP (AB1565, 1:4000, Millipore), β-Tubulin (CW0098M, 1:5000, ComWin Biotechnology), at 4°C overnight. Following washing, the membrane was incubated with horseradish peroxidase-conjugated secondary antibodies (1:10000, ZhongShan Goldenbridge Biotechnology); membranes were revealed with ECL (WBKLS0500, Millipore) and exposed on Luminescent Imaging (Tanon 6200 Luminescent Imaging Workstation, Tanon). Quantification was performed using ImageJ software, and all signals were normalized within the same membrane to β-Tubulin.

### Stereotaxic cannulation and microinjection

At 70 days of age, mandarin voles in the PD group were anesthetized by isoflurane, and 26-gauge bilateral steel guide cannulae (R.W.D. Life Science, Shenzhen, China) were stereotaxically implanted aimed at the PLC (coordinates from bregma: anterior, 2.2 mm; bilateral, ±0.5 mm; ventral, 2.2 mm). Voles were allowed 3 d of post-operative recovery. The CSF (200 nl/side), CSF containing OT (1 ng/200 nl/side or 10 ng/200 nl/side), or CSF containing OT (male 10 ng OT; female: 1 ng OT) plus OTA (10 ng/200 nl/side or 100 ng/200 nl/side) were injected into the PLC. All microinjections were made with a 33-gauge needle that extended 1 mm below the guide cannula and delivered at a rate of 0.1 *μ*l/min as previously described (Young et al., 2014). Data were excluded if tips were located in other brain regions. The final size of each group was 6 except for the 10 ng OT/100 ng OTA male group which numbered 5, and the 1 ng OT/10 ng OTA and 1 ng OT/100 ng OTA female groups (both n = 5).

### Data analysis

Parametric tests were used because all data were normally distributed according to one-sample Kolmogorov–Smirnov tests. Independent sample *t*-tests were used to assess differences in behavior in the OFT (Experiment 1), levels of OTR, V1aR and AVP protein in different brain regions. The significance level was set at *p* < 0.05. The SPT (factors: treatment × stimulus); serum OT and CORT concentration (factors: treatment × sex); mPFC (IOD), NAcc (IOD) and PVN (the number of positive neurons) OT-IR (factors: treatment × sex) were analyzed using two-way ANOVA, paired t-tests were used to compare approach/avoidance with Bonferroni correction for multiple comparisons (given that two groups were examined in Experiment 1, and five groups were examined in Experiment 4, the threshold for significance was set as *p* < 0.025 and *p* < 0.01, respectively). The OFT (Experiment 4) were analyzed using one-way ANOVA. Data were presented as mean ± SEM, all statistical procedures were performed using SPSS 17.0.

## Acknowledgements

This work was supported by the National Natural Science Foundation of China (grants 31372213 and 31170377) and Fundamental Research Funds for Central University (GK201305009). Prof. Tai designed the study and wrote the protocol. Zhixiong He conducted the majority of the work, the statistical analysis and wrote the first draft of the manuscript. Limin Wang, Wei Yuan and Rui Jia discussed the results and provided constructive comments. Wenjuan Hou, Jinfeng Yang, Yang Yang and Luo Luo helped care for voles.

## Additional Information

### Funding

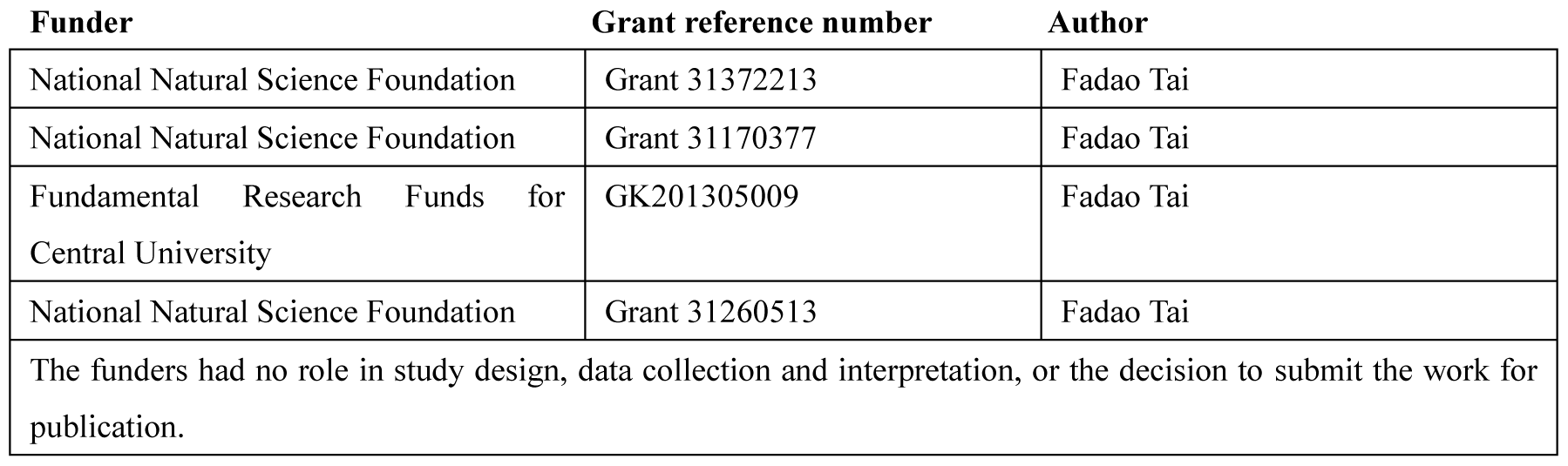

### Author contributions

ZH, Conceptualization, Data curation, Methodology, Writing—original draft, Writing—review and editing; LW, LL, RJ, WY, Conceptualization, Investigation, Methodology, Writing—review and editing; WH, JY, YY, Conceptualization, Resources, Supervision, Methodology, Writing—review and editing; FT, Conceptualization, Resources, Formal analysis, Supervision, Funding acquisition, Methodology, Writing—original draft, Writing—review and editing

### Author ORCIDs

Fadao Tai, <u>http://orcid.org/0000-0002-6804-4179</u>

### Ethics

Animal experimentation: All procedures were approved by the Animal Care and Use Committee of Shaanxi Normal University and were in accordance with the Guide for the Care and Use of Laboratory Animals of China.

---

**Figure 1 - supplement 1.**
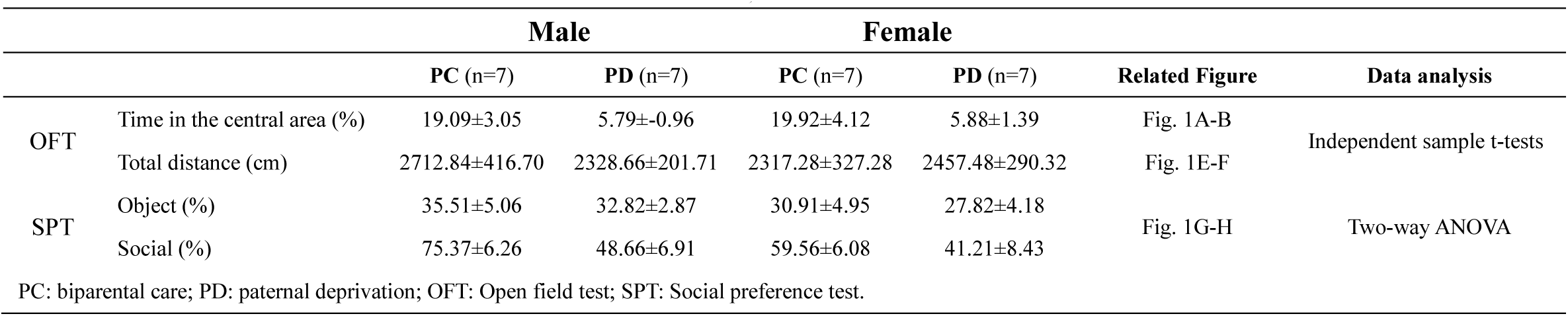
Table 1. Summary of the OFT and SPT analysis.

**Figure 2–4-supplement 2.**
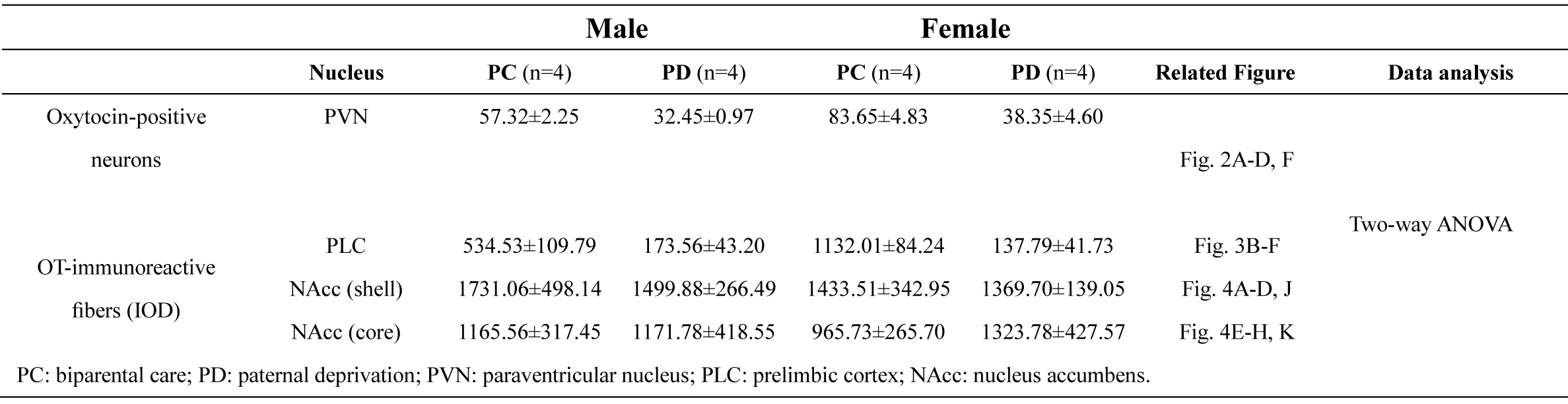
Table 2. Summary of the OT-IR cells or fibers analysis.

**Figure 5-supplement 3.**
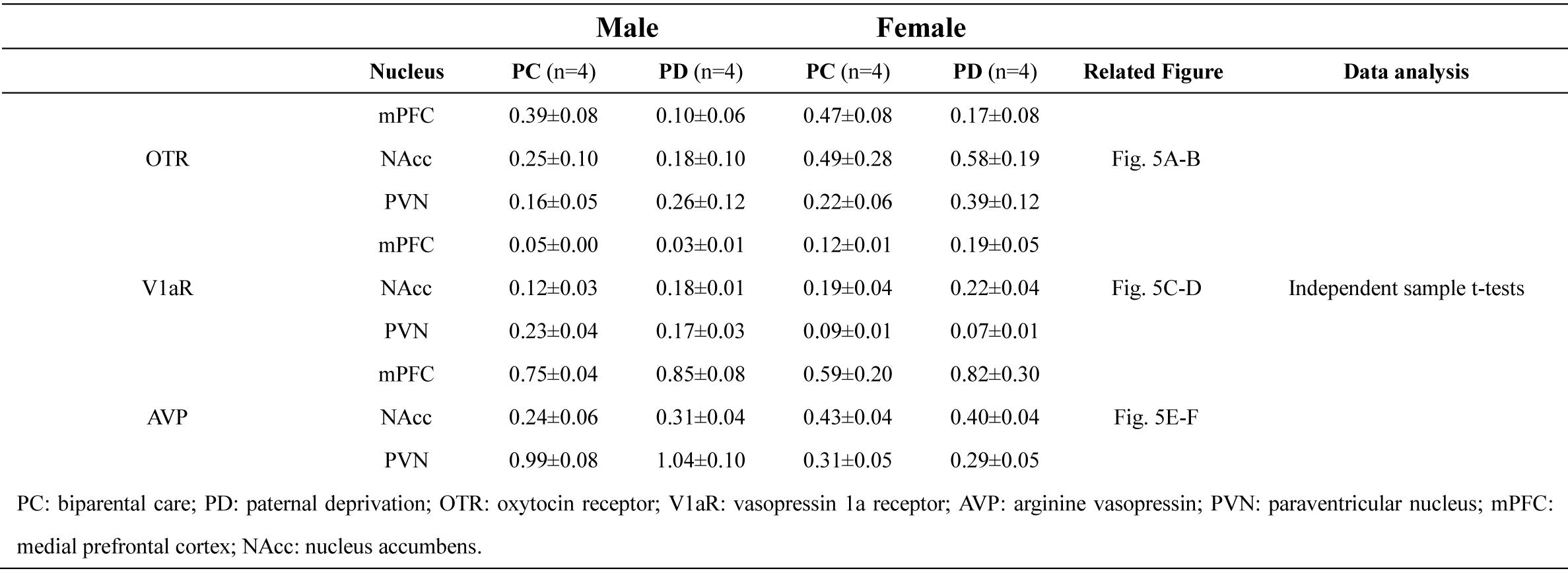
Table 3. Summary of the expression of OTR, V1aR and AVP proteins analysis.

**Figure 6-supplement 4.**
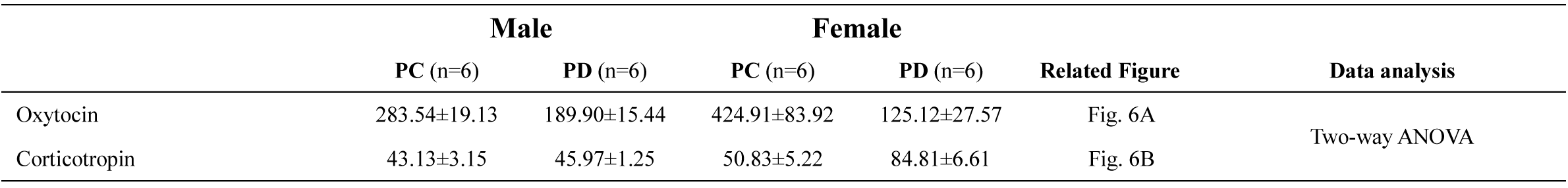
Table 4. Summary of serum oxytocin and corticotropin concentrations analysis.

**Figure 7-supplement 5.**
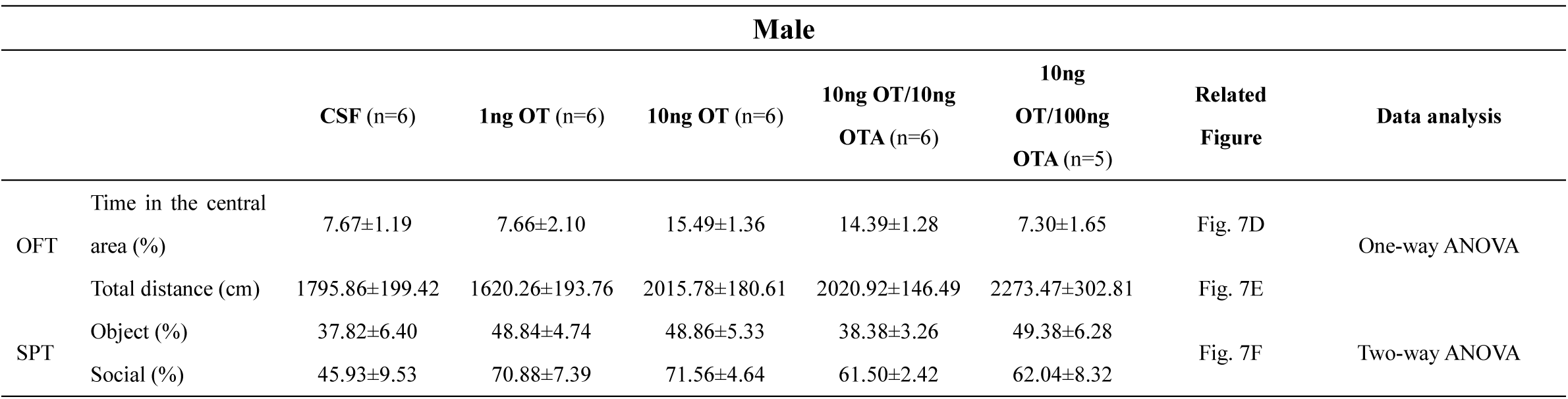
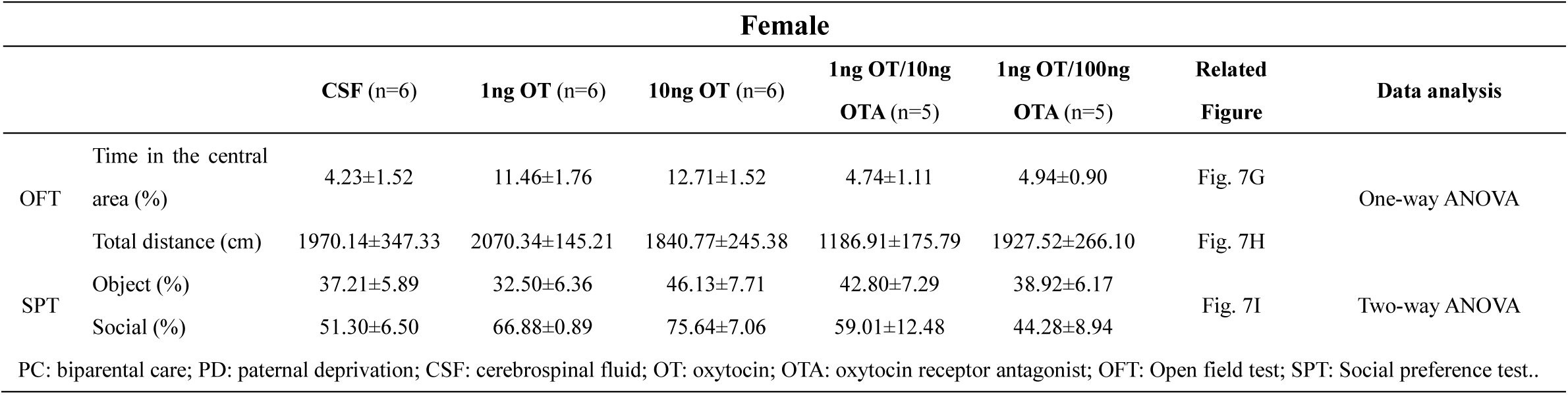
Table 5 Summary of microinjection of OT into the PLC on anxiety-like behavior and social preference analysis.

